# Impact of Climate, Habitat and Scale on the Population Dynamics of Feral Goats on the Isle of Rùm, NW Scotland

**DOI:** 10.1101/2024.06.20.599948

**Authors:** R.I.M. Dunbar

## Abstract

Although feral goats are an invasive species renowned for their ability to survive in degraded habitats, their capacity to occupy high latitude habitats is severely restricted. I analyse long term data on the lifehistory and demography of a feral goat population on the Isle of Rùm, NW Scotland, in relation to both longterm variation in climatic variables and within-population variation in environmental variables. While exhibiting many features characteristic of ungulate lifehistory, goats are especially sensitive to variations in thermal conditions, especially during winter. This is compounded by the fact that, at the latitude of Rùm, goats give birth in mid-winter, even though this imposes significant stress on both mother and kid. Longterm patterns in population growth rates are correlated with winter temperature and the NAO index, with little evidence for density-dependent effects (except in respect of fertility). In addition, there was evidence that the presence of a large sympatric red deer population was limiting the goats’ capacity to increase by denying them access to preferred foraging habitat. Nonetheless, their unusual sensitivity to the thermal environment implies that the goat population will increase significantly with progressive climate warming.

## Introduction

The population dynamics of temperate ungulate populations have been studied in some detail (e.g. Clutton-Brock et al. 1982, 1992), implicating both extrinsic factors (climate, resource quality) and intrinsic factors (density-dependence) as important influences (Milner-Gulland 1994; Post et al. 1997; Post & Stenseth 1999; Coulson et al. 2000; Weladji & Holand 2006; Morellet et al. 2011; Pape & Löffler 2015). However, almost all these studies have focussed on grazers. By comparison, we know a great deal less about browsers. Feral goats (*Capra hircus*) represent one of the most widespread species of browsing ungulates, having successfully adapted to both tropical and temperate habitats on all continents as well as many oceanic islands. Given the significance of this species in both economic and conservation terms (Gould 1979; Hamann 1993; Yocom 1967; King 1992; Dawson & Ellis 1996; Pople et al. 1996; Mitchell et al. 1987; Parkes 1993; Norton 1995; Stronge et al. 1997), our knowledge of its population dynamics is surprisingly poor. In addition, what little we know of this species is confined to tropical island populations, and there have been no published studies of the demography of high latitude populations.

Goats differ from most herding ungulates, such as deer, in having distinct heft groups (relatively stable groups that are attached to a specific territory) (Gordon et al. 1987; Stanley & Dunbar 2013). Within-habitat variation in resource quality and environmental conditions has been identified as playing a potentially important role in the population dynamics as well as community ecology of a number of well studied species (Wiens et al. 1993; Sutherland 1996; Hanski 1999; Thomas & Kunin 1999). Winter survival in red deer (*Cervus elaphus*), for example, has been shown to reflect local population density (Coulson et al. 1997), while marked differences in various lifehistory parameters have been noted between adjacent Soay sheep (*Ovis aries*) heft groups (possibly reflecting micro-habitat differences in grazing opportunities: Coulson et al. 1999). Similarly, Lee & Hauser (1998) found that differences in birth and survival rates between neighbouring groups of vervet monkeys (*Cercopithecus aethiops*) correlated with differences in local resource availability.

Here, I use data from a population of individually known feral goats inhabiting the Isle of Rùm, NW Scotland, to explore (a) the impact of micro-habitat variation in key climatic and resource variables on lifehistory processes at the local scale and (b) the influence of climate and population density on long term demographic trends at the population level.

## Methods

The study was carried out on the west coast of the Isle of Rùm NNR (latitude 57°N), a large island lying off the northwest coast of Scotland. The main study population consisted of all the goats resident in a 10-km stretch of sea cliffs bordering the western side of the island between Harris Bay in the south and Glen Guirdhil in the north, representing about half the island’s total goat population. Goats were known to have been feral on the island as early as the 1770s, though most of the present population are the descendants of animals abandoned by the island’s crofters when these were persuaded to emigrate to Canada in the 1820s (Gordon et al. 1987). Studies of this population were carried out in 1980-2, 1993 and 2000-2006.

The study area is dominated by high sea cliffs rising 200-400m above sea level above a rocky shoreline platform. In its upper reaches, it gives rise to shallower scree-covered slopes rising to bare, stoney, cliff-bound mountain tops at ∼600m asl. The northern and southern boundaries are defined by 1km-deep grass- and heath-covered valleys (Glen Guirdhil and Glen Harris, respectively) that slope gently inland to the base of the island’s rugged central mountains that rise abruptly to altitudes of ∼700m.

Rùm experiences an Atlantic seaboard climate. Fig. 1 shows mean monthly rainfall and mean monthly temperatures for Harris Bay at the southern edge of the study area. Annual rainfall on Rùm averaged 1513 (±209) mm over a 13-year period (1969-1981 inclusive). The period July-January tends to receive more rain than other months, but the pattern is very variable. Temperatures are lowest between December and April, though minimum temperatures rarely drop below freezing by more than a 2-3°C or for more than a few days each month. Winter gales are common, with high wind strengths adding a significant degree of windchill to minimum temperatures on the higher ground throughout the year.

**Fig. 1.**
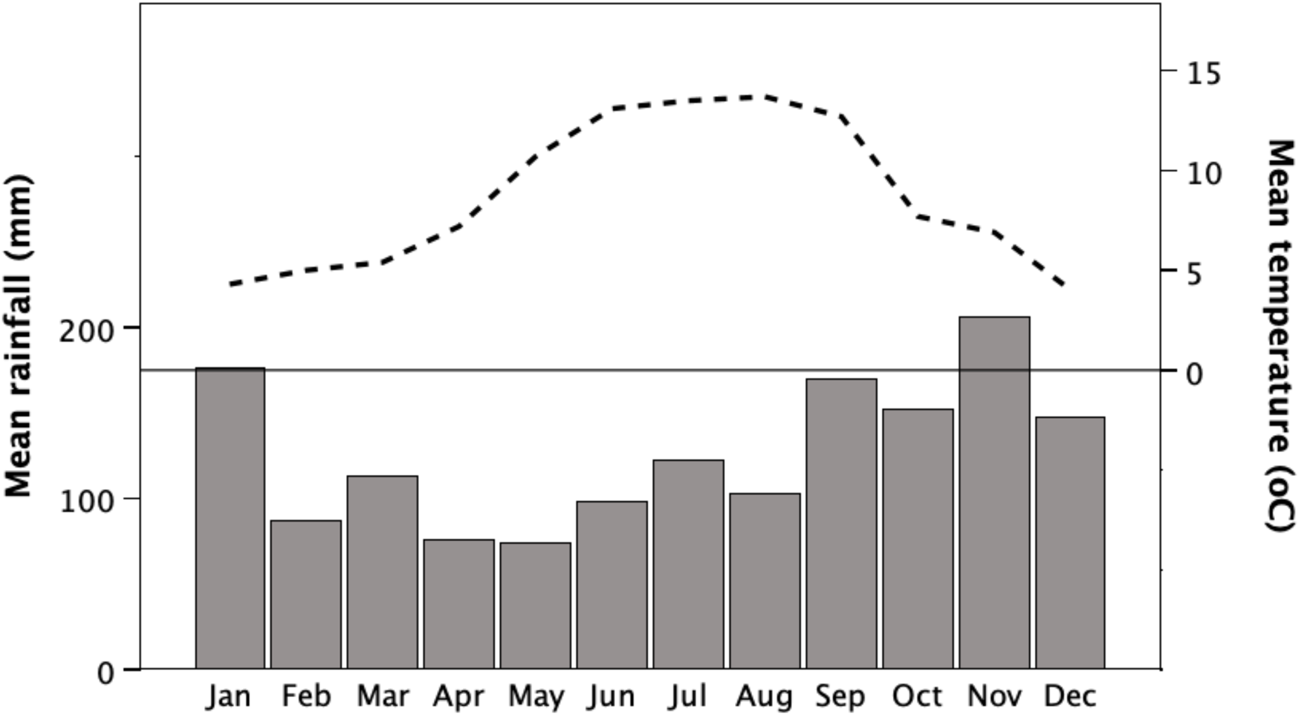
Climate at Harris, Isle of Rùm. Bars (lower panel): mean monthly rainfall (mm) for 1969-1981. Dashed line (upper panel): mean daily temperature by month for 1981-1982. Horizontal line defines 0°C.

In order to assess thermal variability across the habitat, min/max thermometers were set at a number of sites that varied both in altitude up the cliff face and in degree of shelter (including the rear wall of two caves at beach level). These included sheltered sites on the open beach at Harris and Wreck Bay (below the Sgorr Reidh cliffs), at the rear of two caves at beach level in Wreck Bay, half way up (100m asl) and at the top (200m asl) of the Sgorr Reidh cliff above Wreck Bay, and on the top of Bloodstone Hill (388m asl). Thermometers were read at the beginning of each month. The distribution of monthly minimum temperatures (Fig. 2) shows that there is a 3°C thermal gradient up-slope from beach level to the highest point in the study area (Bloodstone Hill); in addition, there were significant thermal gains to be had from using the beach level caves, with the shallow cave offering an advantage of about 1.5°C over the open beach and the deep cave about 3°C. These values are reflected in the frequencies with which temperatures fell below freezing. Monthly minimum temperatures never fell below freezing in the deep cave, but did so in 12% of months for the shallow cave, 31% for the beach (two locations), and 56% at the top of the cliffs and 59% on Bloodstone Hill (*N*=16-18 months each).

**Fig. 2.**
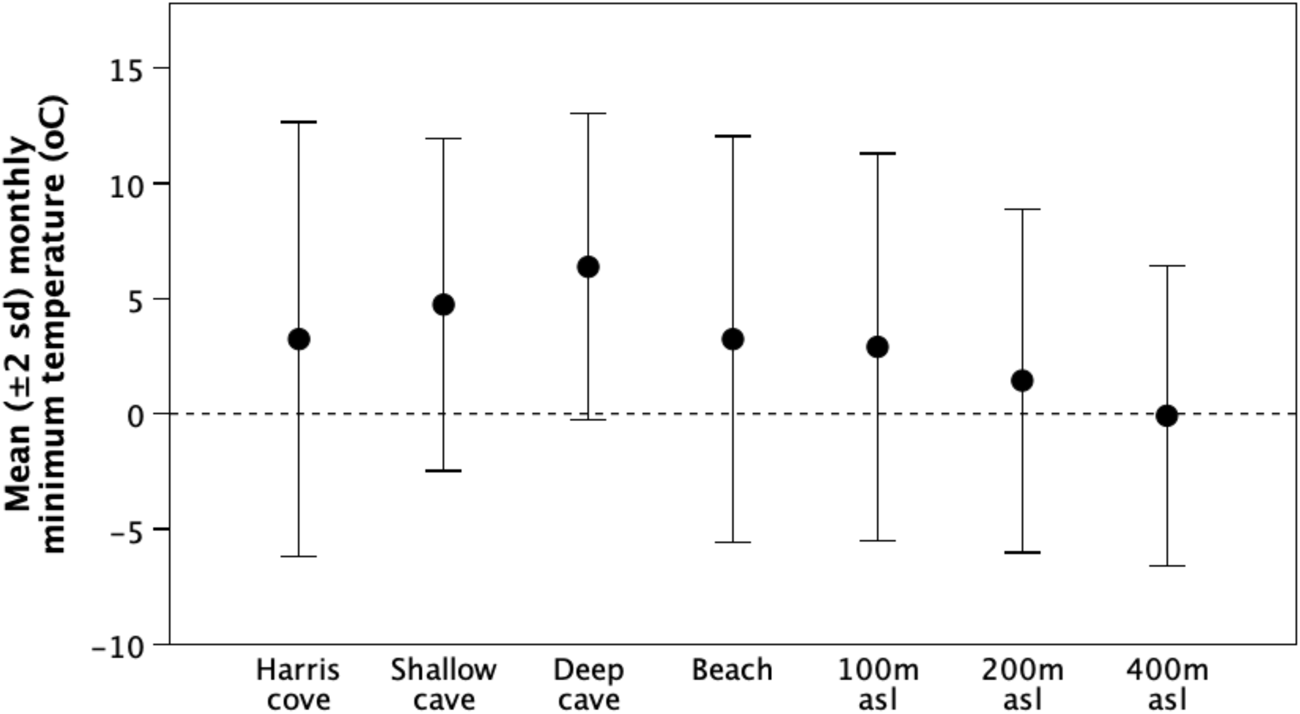
Mean±2sd monthly minimum temperature (1981-1982) at different locations on the west coast of Rùm. Locations differ by altitude: beach (2m asl), mid-cliff Wreck Bay (100m), Sgorr Reidh cliff top (200m), Bloodstone Hill (400m). The two caves differ considerably in depth.

Long term rainfall and temperature for Tiree (60 km southwest of the study site) were sourced from the UK Meteorological Office (https://www.met-office-historic-data.co.uk/chart), supplemented by data from the island’s reserve office for Harris itself. Data for the North Atlantic Oscillation (NAO) index were obtained from Hurrell (2002). NAO indexes the air pressure difference between the subtropical (Azores) and sub-polar (Iceland) eastern Atlantic, which determines whether the northwest coast of Europe experiences warm tropical or cold polar conditions: high positive values predict warm wet, while negative values predict cold dry conditions (Hurrell 1995). A number of studies have shown that the NAO index correlates with the population dynamics of northern European red deer populations: high winter NAO indices associated with warm, wet winters are correlated with high ungulate mortality (Forchhammer et al. 1998).

The vegetation on the west coast of Rùm is a mosaic of *Calluna*/*Erica* heath and a variety of water-logged habitats (wet heath, blanket bog, *Molinia* fen and *Schoenus* fen), with *Festucca/Agrostis* sward on the better drained flat areas at sea level and on the two valley floors. In 1980, the vegetation was sampled along 19 transects, each of 100 sample points located 1.2m apart, set at right angles to the cliff line at regular intervals across the main goat habitat; the presence of a plant species was recorded for each point if vegetation of that species touched the pin placed vertically at the sample point. These data were used to calculate the proportion of all sample points at which a given type of vegetation was present. Vegetation growth was recorded in two exclusion plots in 2000. Biomass was at its maximum during May (heath habitats) and June (sward habitats) and had declined sharply by August (see also Gordon 1989a). Crude protein (g/100g dry matter) and energy (MJ/kg dry matter) of the main forage vegetation (heaths, grasses and sedges) were determined from monthly samples (analyses carried out by the UK government’s ADAS laboratory, Wolverhampton). Both nutrients were at their lowest in November/December, increased steadily through the spring and summer and reached their maximum in June/July.

Every goat in the population was individually identified by coat colour and horn shape. Except for kids with known birth dates, all goats were aged by counting annual horn rings. Although annual horn rings can be harder to distinguish in older animals (Bullock and Pickering1984), ageing by horn rings is completely accurate in younger animals and reliable in older ones (for further details, see Dunbar et al. 1990). Females were permanent residents of one of half a dozen heft groups whose ranges fell wholly within the study area. Males were more mobile, often spending extended periods of time with neighbouring populations (Gordon et al. 1987). At the latitudes of the British Isles, goats do not forage at night (Shi et al. 2003; Dunbar & Shi 2013), but instead rest up in caves and other sheltered locations at sea level. A complete survey of all caves along the beach was undertaken in 1981.

An individual was assumed to be alive provided it was seen at least once a week in the study area. Any female or kid that was missing from the records for more than two weeks was presumed to have died. Males were assumed to have died in the week after they were last seen if they did not subsequently reappear in the study area or were not located elsewhere during periodic searches outside the main study area. Kids were recorded as having been born in the week when they were first seen. If a female suspected to be pregnant went missing, a particular effort was made to search the lower cliff lines within her home range since most births occurred on or near the beach platform and kids were generally kept low during the first few months of life. It was not always possible to sex kids at a distance until they were 6 months old: for kids that died before they were old enough to sex, I have assumed an even sex ratio since those kids that were sexed were of approximately even sex ratio.

Estimates of the total population size have been obtained in a number of years since 1960 by NatureScot staff carrying out complete surveys of the entire west coast of the island between Dibidil on the southern tip and Kilmory in the north. This encompasses the entire area occupied by goats on Rùm. The survey was normally undertaken during the late spring, with teams of observers covering different sections of the island. Although it is likely that some animals were missed, census effort was reasonably constant and the figures are close to those obtained from our more detailed counts of individually identified animals. Censuses were carried out in 17 individual years between 1960 and 1978, but were intermittent thereafter. Estimates of population size are available for 12 pairs of consecutive years; in a number of additional instances, censuses were taken two years apart and in these cases the annual growth rate was estimated from the standard Lotka-Volterra equation assuming a constant growth rate across both years. To these we can add the growth rate within the Harris sub-population during our study.

## Results

### Mortality rates

Table 1 gives the age-specific standing crop for each sex at the beginning of the three main study periods. Sex-specific survivorship curves for females and males constructed from age-specific death rates are shown in Fig. 3. Separate indices are shown for the three study periods for females, but to avoid cluttering the graph too much only a mean value for 1981-2 and 2000-1 is shown for males. No animal in the population was older than 12 years. Males had somewhat lower mortality rates than females of the same cohort in their early years, but thereafter their mortality rates increased considerably, mainly because of the costs involved in rutting. For males, the rut is intense from the yearling stage onwards, but hits a peak between the ages of 4-6 years (Dunbar et al. 1990, Lloyd 2003). By contrast, female mortality is relatively constant at 10-20% per annum up to age 9 years, after which it rises steeply. The final point to note is that female survival was much higher in 2003-4 than in either of the earlier study periods.

**Fig. 3.**
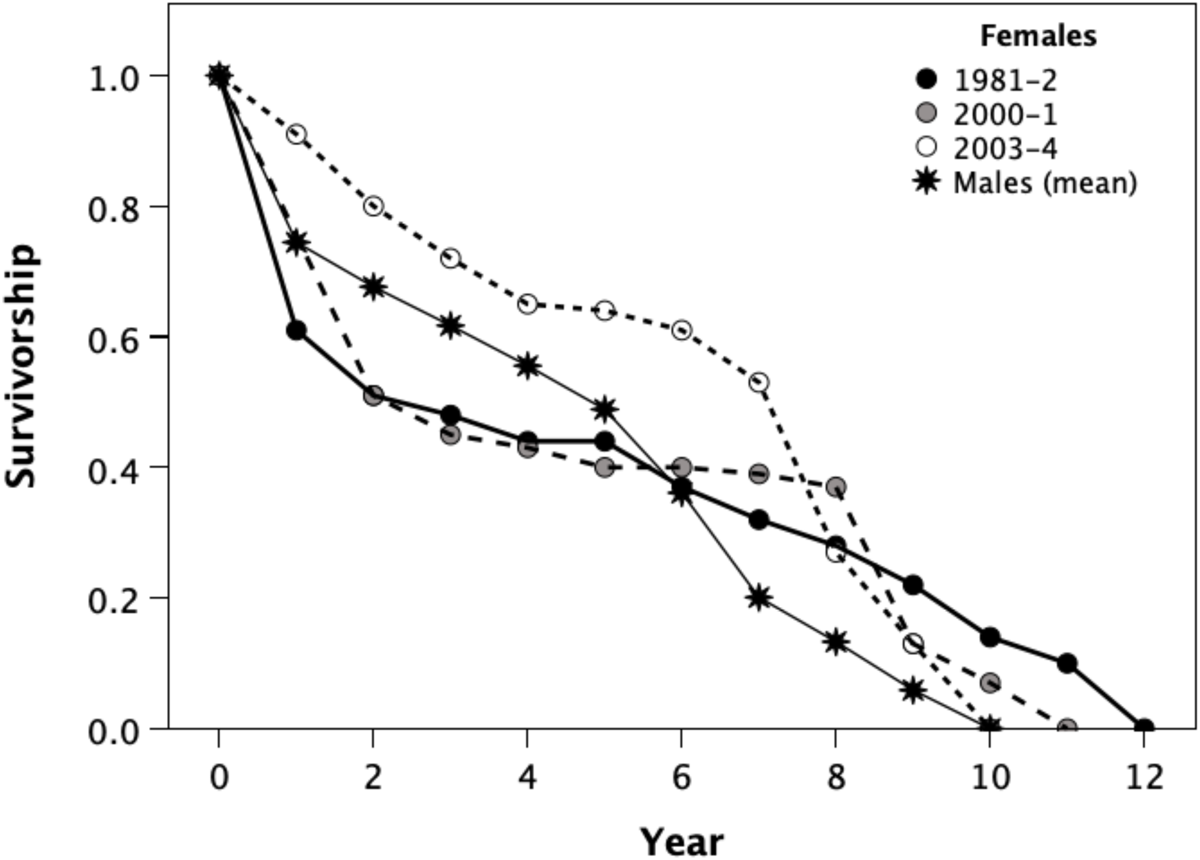
Survivorship curves for female goats in three separate study years (1981-2, 2000-1, 2003-4) and mean survivorship for males (1981-2 and 2000-1).

**Table 1.** Population age structure in individual study years.

| Year | 1981 |  | 1982 |  | 2000 |  | 2001 |  | 2002 |  |
| --- | --- | --- | --- | --- | --- | --- | --- | --- | --- | --- |
| class§ | ♂ | ♀ | ♂ | ♀ | ♂ | ♀ | ♂ | ♀ | ♂ | ♀ |
| 0* | 10 | 9 | 6.5 | 5.5 | 8 | 12 | 6 | 11 | - | - |
| 1 | 7 | 6 | 11 | 5 | 5 | 14 | 6 | 10 | 4 | 7 |
| 2 | 7 | 10 | 7 | 6 | 12 | 7 | 4 | 14 | 4 | 5 |
| 3 | 9 | 6 | 8 | 9 | 7 | 15 | 11 | 6 | 3 | 12 |
| 4 | 7 | 2 | 10 | 6 | 18 | 6 | 7 | 14 | 10 | 6 |
| 5 | 2 | 4 | 8 | 2 | 9 | 3 | 14 | 6 | 5 | 14 |
| 6 | 4 | 4 | 2 | 4 | 16 | 16 | 8 | 3 | 14 | 4 |
| 7 | 1 | 3 | 1 | 4 | 3 | 2 | 14 | 15 | 8 | 1 |
| 8 | 2 | 3 | 2 | 3 | 2 | 3 | 3 | 2 | 14 | 11 |
| 9 | 0 | 1 | 1 | 2 | 0 | 1 | 2 | 2 | 1 | 0 |
| 10 | 0 | 3 | 0 | 1 | 0 | 1 | 1 | 1 | 2 | 0 |
| 11 | 0 | 1 | 0 | 2 | 1 | 0 | 0 | 1 | 1 | 0 |
| 12 | 0 | 0 | 0 | 0 | 0 | 0 | 1 | 0 | 0 | 1 |
| Total | 51 | 52 | 56.5 | 49.5 | 81 | 80 | 77 | 85 | 66 | 61 |

§ ♂=males; ♀= females
\* kids born in January that died before 1<sup>st</sup> February are not included
Source for 2000-2002: Lloyd (2003)

Male and female deaths were distributed differently across the year (Fig. 4). Male mortality clusters very strikingly in the autumn months, reflecting the energetic costs of the rut during August and September. Male feeding rates are drastically reduced during the rut when their attention is focussed almost exclusively on the business of mating (see Dunbar & Shi 2013). The rut involves intense fighting between males, and those who have been very active in the rut hit a combination of deteriorating forage conditions in the autumn and worsening weather in an already weakened condition. Examination of carcasses revealed that most males were emaciated when they died, and many were suffering from conditions (e.g. foreleg lameness, sand-cracking of hooves) that would have seriously hampered their ability to forage effectively.

**Fig. 4.**
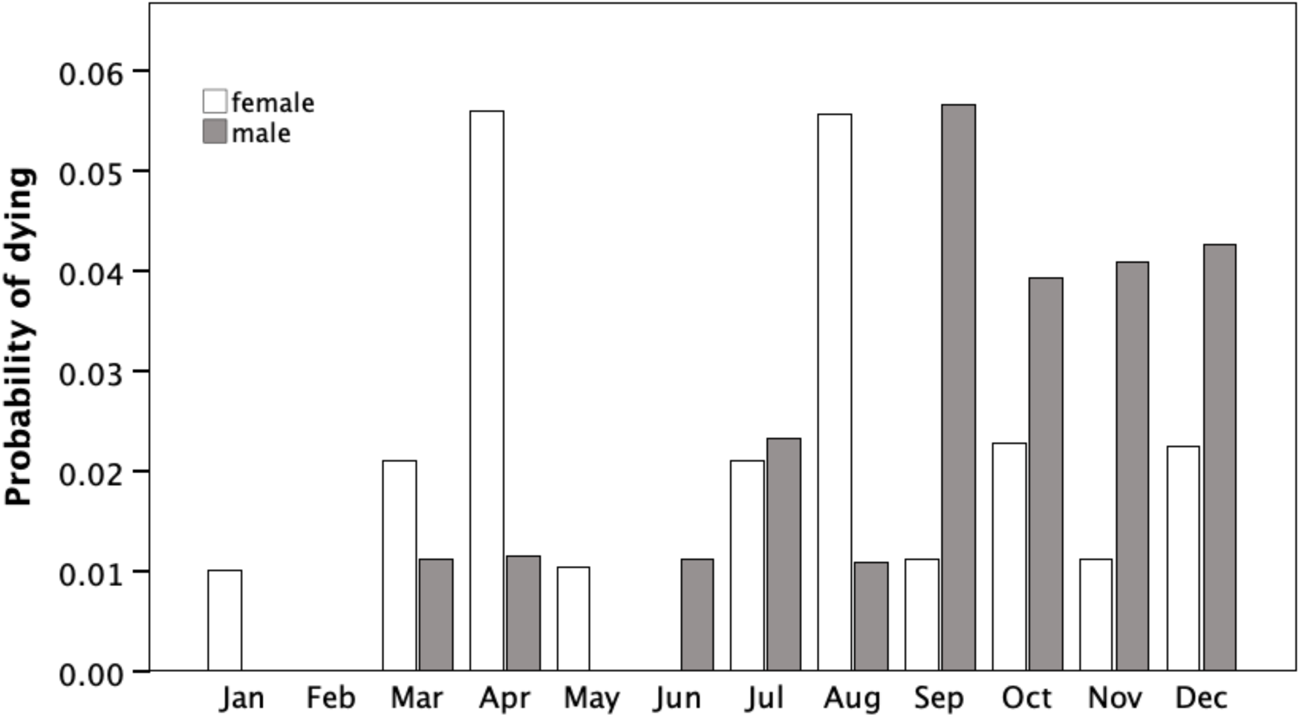
Mortality in adult males and females (>6 months of age) by month.

Female deaths were more evenly distributed than those of the males, but tended to cluster during the late spring and summer (Fig. 4). This mainly reflects the costs of lactation under conditions that are energetically stressful (Dunbar et al., submitted). The fact that mortality is at its peak during the summer when vegetation productivity is at its highest (Gordon 1989a) suggests that reproduction can tax older females to a point where they cannot recover despite improved forage conditions.

Comparison of heft-specific female mortality rates for the 1981-2 study with contemporary estimates of the density of six different vegetation types (grasses, *Molinia* spp., *Eriophorum* spp., sedges, forbs, heaths) and the number of caves (used as nighttime shelters) indicates that within-heft adult female mortality is related (negatively) only to the number of caves (Fig. 5: Spearman *r_s_*=-0.90, N=5, p=0.037) and the number of females in the heft (Spearman *rs*=1.00, N=5, p=0.037, pointing, in the first case, to the costs of thermoregulation during periods of low temperature especially at night as the principal factor influencing differences in female survival at the heft level and, in the second case, to some evidence for density-dependent mortality. Because males are less strongly attached to the hefts they were born into, and roam more widely, heft-based mortality rates are less meaningful in their case.

**Fig. 5.**
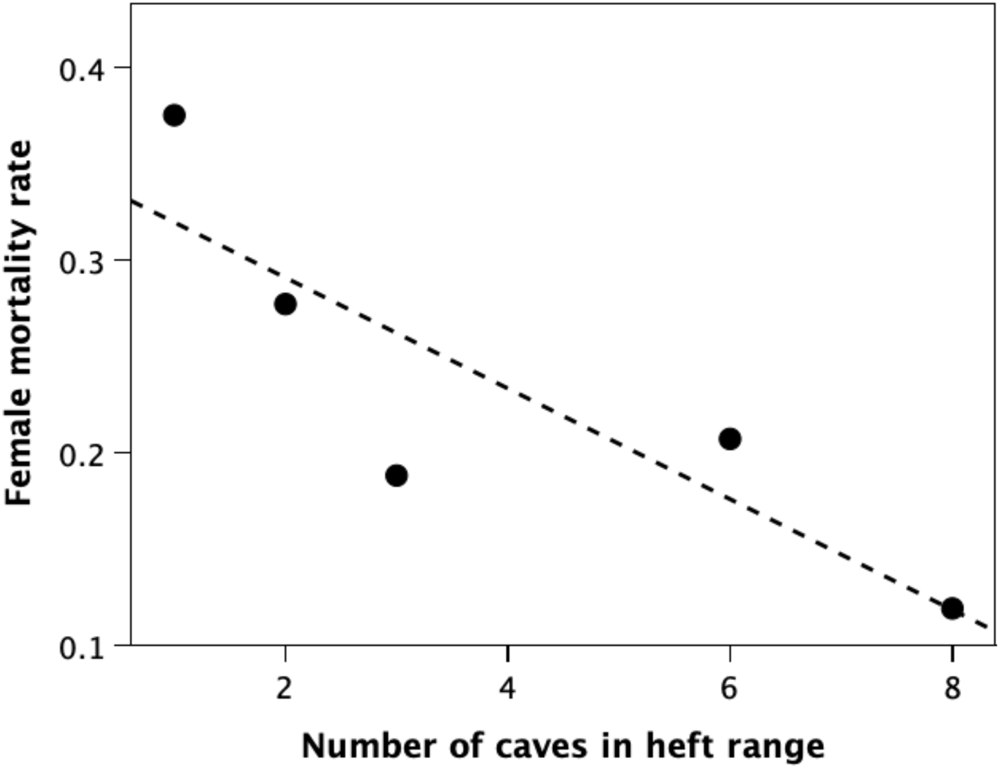
Adult female (>6 months old) mortality rates for each of the five heft groups, plotted against the number of caves (used as nighttime shelters) in the heft’s range.

### Fecundity

Mean birth rates for females in each year cohort are given for the three main study periods in Fig. 6. Fecundity exhibits the classic mammalian hump-shaped pattern, with the likelihood of giving birth at its maximum in mid-reproductive career (between 3-6 years of age, in this case). Within this general pattern however, fecundity rates vary considerably, with rates being highest in 2003/4 and lowest in 1981/2. Overall, birth rates are quite modest, with most females giving birth to singletons only in alternate years between ages 2 and 9. Of 128 females from two study periods whose reproductive histories could be traced over three successive years (five successive birth peaks), 33 (25.8%) produced no kids at all, 51 (39.8%) produced a single kid, 43 (33.6%) produced two, and one (0.7%) produced three (because her first two kids died). Although twinning is the norm in tropical goat populations (Singh et al. 2009), only one female produced twins in the 10 annual births seasons that this population was under study (290 reproductive age females), and both kids died within a few weeks of birth.

**Fig. 6.**
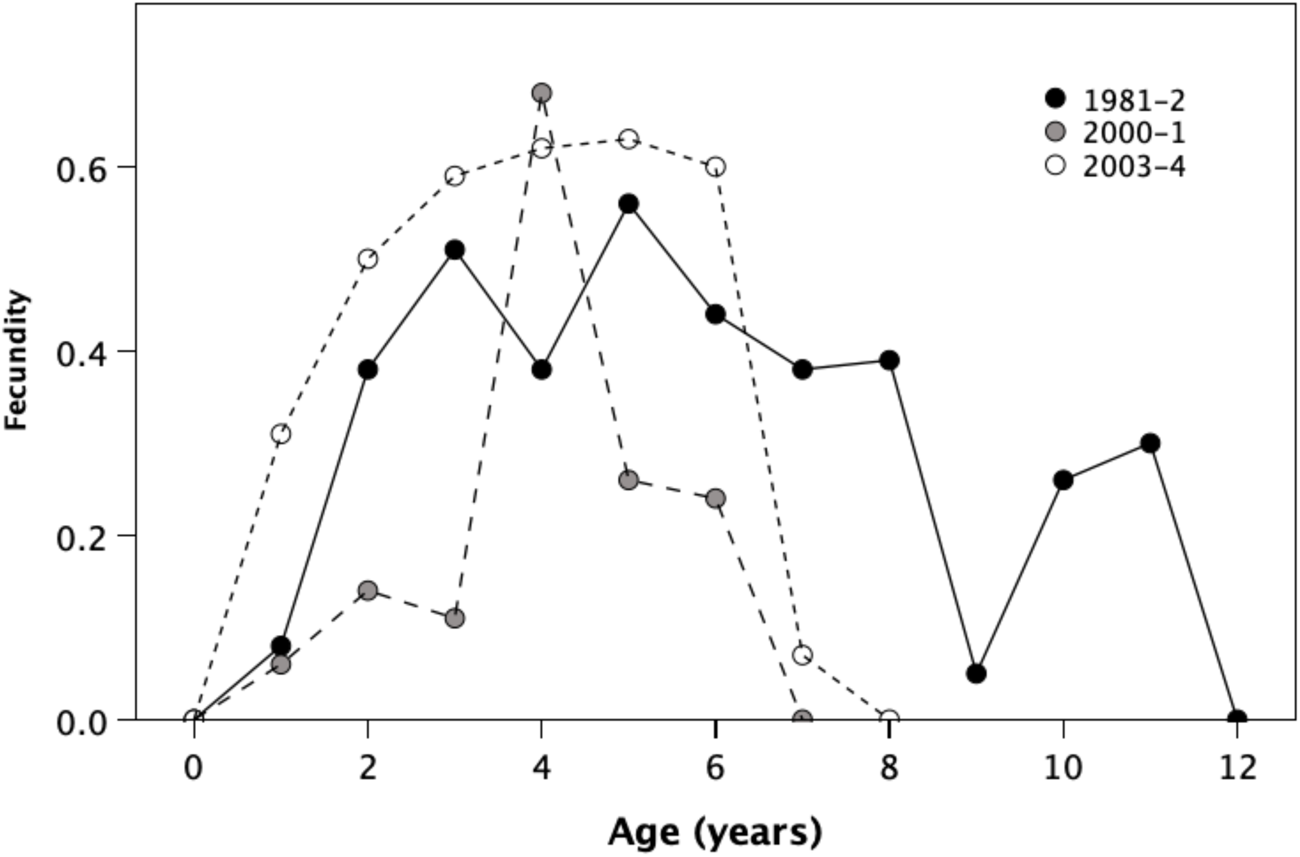
Mean fecundity (births per female per year) plotted by age of female for three study years. The precipitate drop in birth rates at age 5-6 in 2000 and 2003 reflect the impact of the new Harris heft created mainly by younger females. Data for 1981-2 are from GnP, Sgorr Reidh, A’Bhrìdeanach, Camas and Bloodstone hefts; for 2000-1 from Harris, GnP, Sgorr Reidh and A’Bhrìdeanach hefts; for 2003-4 from Harris, GnP and Sgorr Reidh hefts.

Births are highly seasonally distributed (Fig. 7) with 67% of kids being born in late January and February. Paradoxically, these are the months that experience the lowest temperatures and some of the worst weather – reflecting the fact that this subtropical species has not spent sufficient time at these latitudes to adapt fully to the climatic regimes they find themselves in. Females who lose kids within the first few weeks of birth return to oestrus immediately, resulting in a small secondary birth peak in July-August (accounting for 22% of all births).

**Fig. 7.**
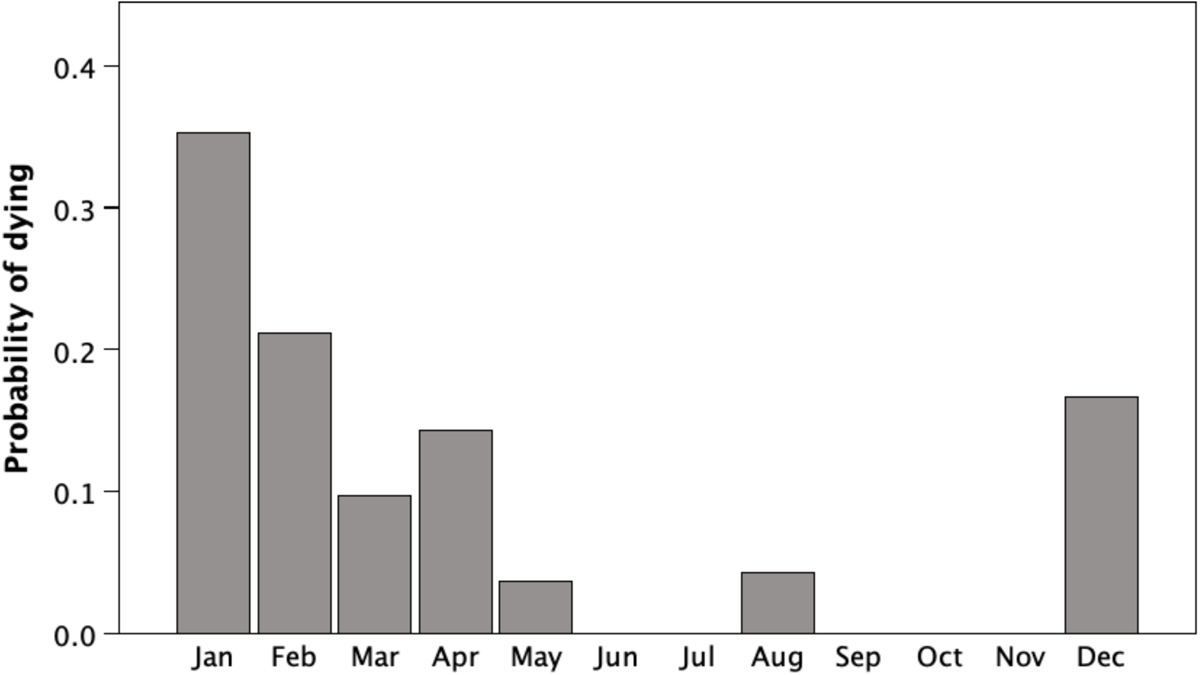
Mean number of kids born per month in 1981 and 1982.

There are marked differences between the hefts, with heft-specific birth rates varying by as much as a factor of two (Fig. 8). Stepwise regression (with the climatic and vegetational indices given in Table 2 as independent variables) suggests that virtually all this variance can be explained by two just features of a heft: the amount of heath vegetation available and the number of females per cave. The best-fit regression equation is:

**Fig. 8.**
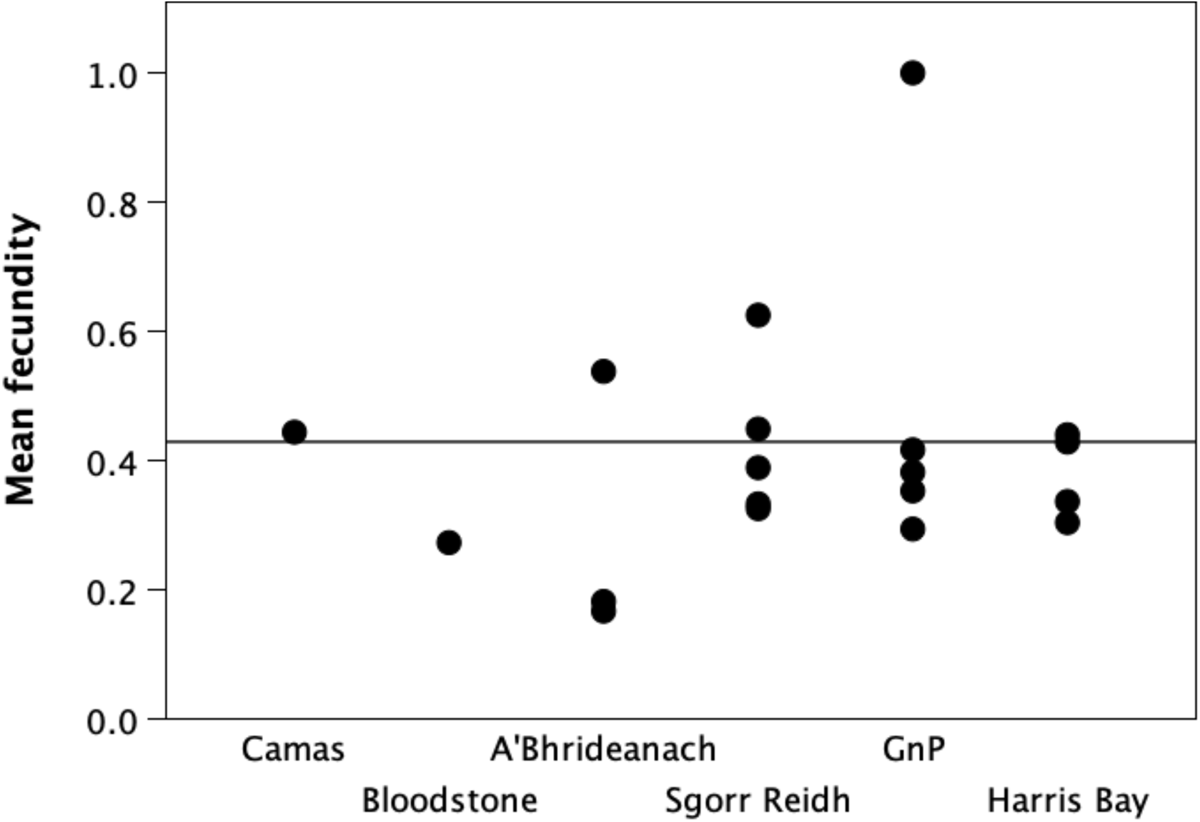
Mean fecundity in each study year for each heft group. The horizontal line demarcates the overall average replacement rate (two surviving offspring in a reproductive lifetime), based on 50% kid mortality (see Figs. 1 and 7) and a 7-year reproductive lifespan (see Figs. 1 and 6). The hefts are listed, left-right, in order north to south. Note that in many years most of the hefts are struggling to maintain demographic stability. Data derive from all three study periods.

**Table 2.**
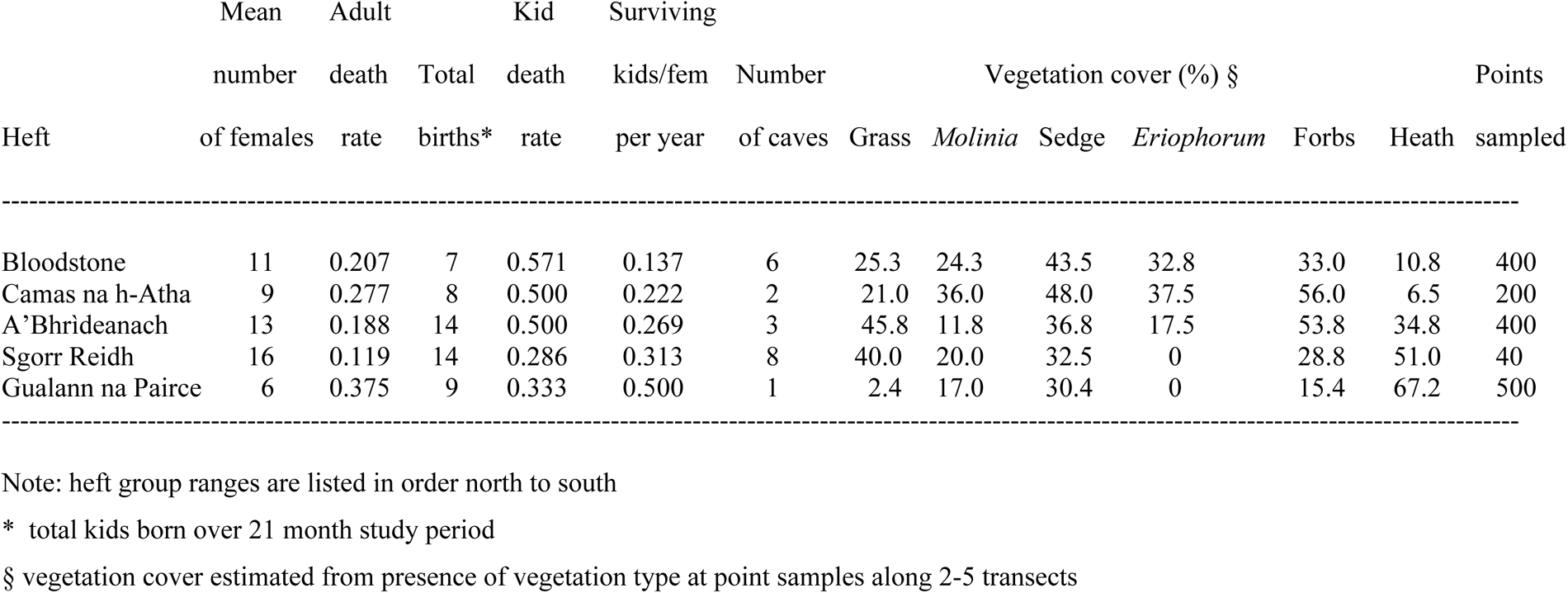
Mortality and fecundity rates for adult females (>6 months old) and mortality rates for kids (<6 months old) in five female heft groups for 1981-2.

Birth rate = 0.159 + 0.0033 (heath cover) + 0.061 (females per cave)

(*r*^2^=0.996, F_2,2_ = 270.2, *P*=0.004; partial regressions: heath cover, *t*=11.40, *P*=0.008; females per cave, *t*=14.50, *P*=0.005).

Note that slope coefficients are both positive. Pregnancy rates were higher when there was more heath cover available, reflecting both the availability of a core food resource (Gordon 1989b) and the fact that heather offers some shelter from wind; in addition, pregnancy rates were higher when more females were packed into each cave at night. That the latter reflects thermoregulatory costs is indicated by the fact that if the number of females in the heft is substituted for the number of females per cave the relationship is no longer significant. The presence of other animals is likely to add significantly to the nighttime temperature within a cave in addition to the effects due to shelter from wind (see Fig. 2).

Fig. 9 plots mortality rates on newborn kids over their first year: 97% of kid mortality occurs in the first month. This is mainly due to the fact that kid mortality during any given quarter is inversely related to the mean ambient temperature (Fig. 10: t=-2.54, p=0.052). Winter-born kids experienced mortality rates that were around three times higher than those for summer-born kids (18 deaths among 49 winter-born kids vs two deaths for 15 summer-born kids). Nonetheless, most kids that make it through to week 9 have a fairly high chance of surviving to sexual maturity.

**Fig. 9.**
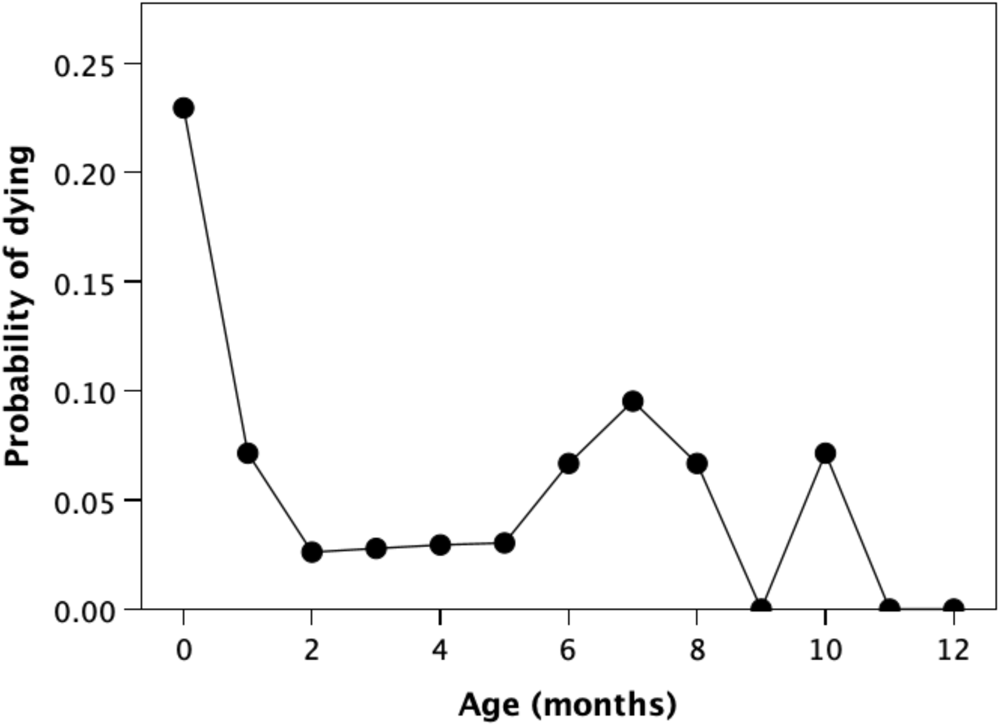
Monthly mortality rates for kids over the first year of life, based on data for 1981-2.

**Fig. 10.**
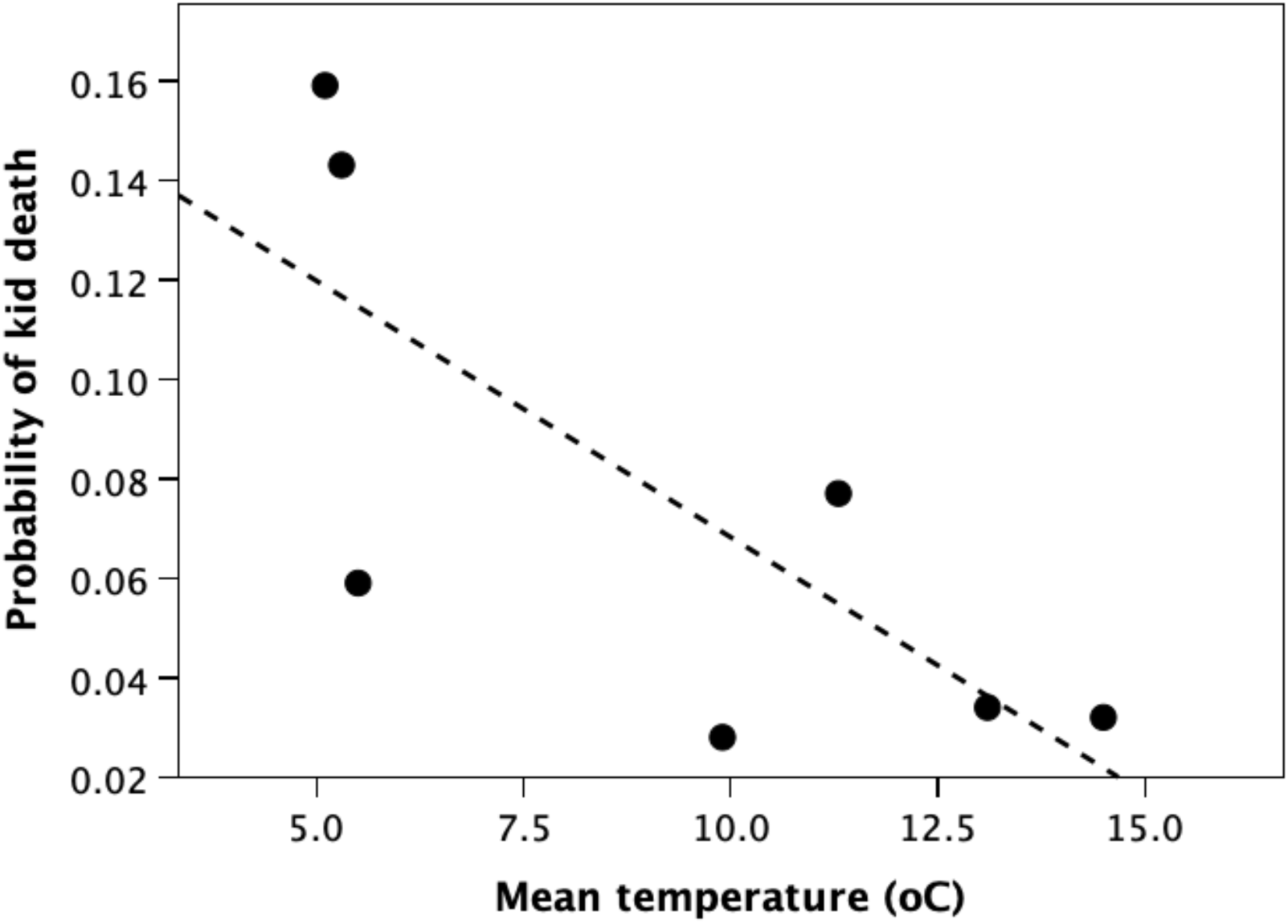
Kid mortality in individual three-monthly time blocks plotted against the mean temperature. Mean temperature is the mean of the monthly means (this being the midpoint between mean daily minimum and mean daily maximum temperatures for that month). Based on data for 1981-2.

Table 2 compares kid mortality rates at the heft level for the 1981-2 sample. This necessarily held ambient temperature conditions constant. Two heft groups (Guallan na Pairce [GnP] and Sgorr Reidh) appeared to have much lower kid mortality rates than the other three. Comparing kid mortality rates against the density of the six main vegetation types and the number of caves, the strongest univariate relationships were a positive correlation with the density of *Eriophorum* (an indicator species for water-logged habitats: t_3_=3.63, p=0.036) and a negative relationship with the density of heath habitats (a major component of the diet: t_3_= -2.86, p=0.065). The availability of caves had no effect, suggesting that the main factor influencing kid mortality may be energy throughput from maternal lactation.

### Population dynamics

Fig. 11 plots the sizes (indexed as the number of females older than 6 months) of the seven heft groups in the main study area over time. In general, heft group size seems to remain relatively stable through the 1980s and 1990s, and then explode from the late 1990s. Notice that the Harris heft group did not exist prior to the late 1990s, and when it does come into existence it increases in size very rapidly. Until the late 1990s, Harris Bay was effectively a goat-free zone (aside from occasional transient males on their way to or from the Papadil area on the south side of the bay). Harris seems to have been colonised by a group of females from the Papadil area to the south, which had a much larger population even in the 1980s (upper left down-triangles in Fig. 11). Their appearance coincides with a rapid increase in all heft group sizes in the late 1990s, suggesting that high densities might have persuaded some females to seek new ranges with fewer females.

**Fig. 11.**
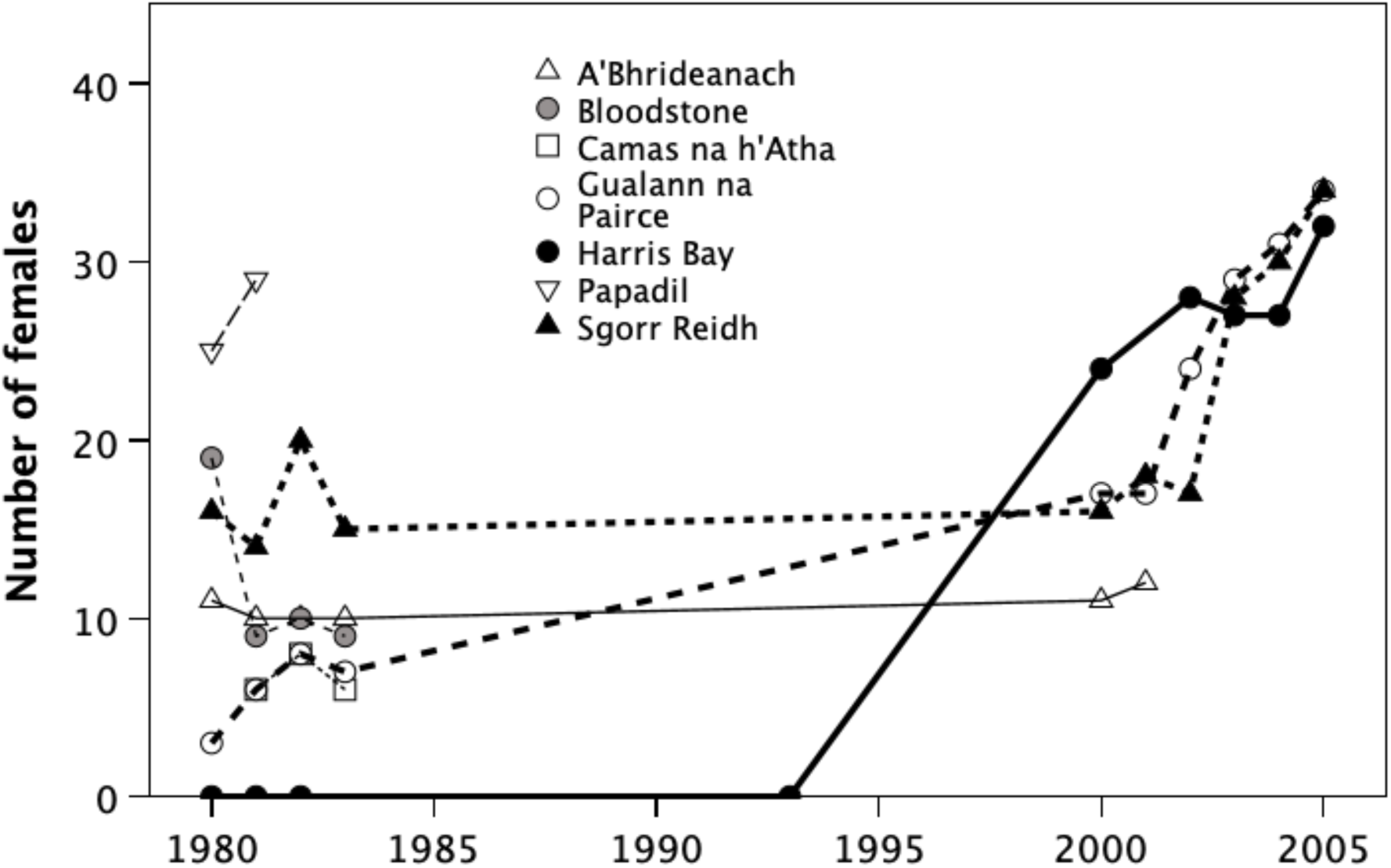
Heft size (number of females >6 months) for individual hefts over time.

There would, thus, seem to be two separate effects to explain. One is the small scale oscillations in individual trajectories present in all hefts across time. The second is the dramatic increase in group size across all hefts beginning in the later 1990s. To explore these, I regressed annual population growth rates from the NatureScot goat census (which provides a much longer time span and a larger sample) against annual rainfall in both the current and the previous year, winter (January-March) NAO index, and population size in the previous year (reflecting the potential impact of density-dependent ecological competition).

Fig. 12a plots goat population size against winter NAO, while Fig. 12b plots the proportional growth rate against NAO (discriminating by decade). Based on a *k*-means cluster analysis of the residuals from the regression line for the lefthand (cold, dry) datapoints, the data partition into three quite separate grades (F_2,19_=109.6, p<<0.0001) corresponding to very different NAO regimes, with growth rates strongly positively correlated with NAO value *within* regimes. This points to a direct effect of temperature on survival. However, there also seems to be a negative trend *across* these NAO regimes (residual regressed on year: standardised β=-0.528, r^2^=0.278, F_1,20_=7.72, p=0.012), suggesting that growth rates become increasingly negative in wetter conditions, causing the peak in growth rates under optimal conditions to fall. It is notable that most the lefthand group in Fig. 12 are early years (1960s) while the right hand ones are mostly much later (1970s and 1980s). In fact, there is a significant correlation between NAO winter index and date (r=0.676, N=27, p<0.0001). This suggests that the climate was getting steadily warmer, as is confirmed by the obvious trend in winter (January-February) minimum temperatures (Fig. 13).

**Fig. 12.**
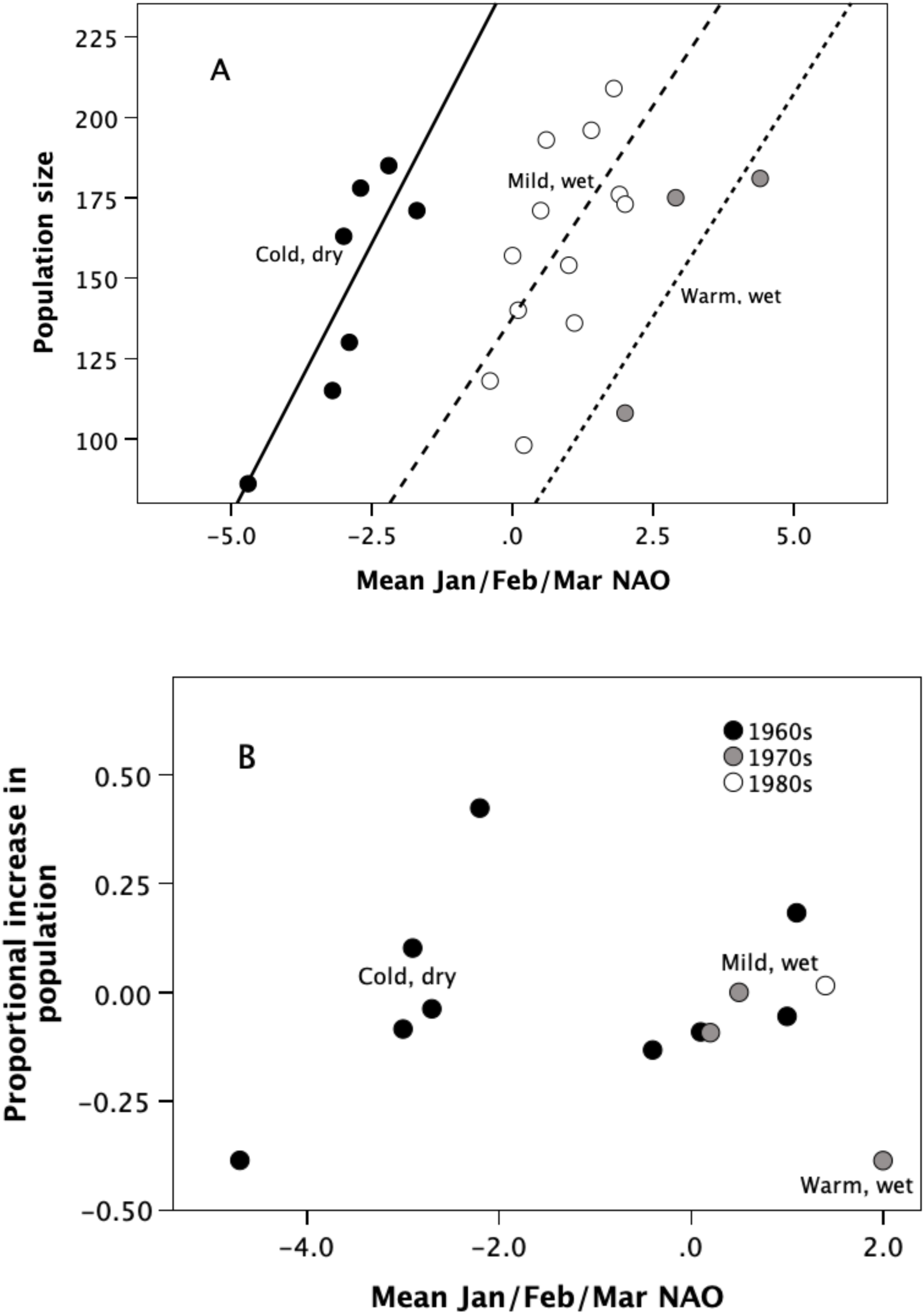
(a) Population size and (b) growth rate (*r*) in each year plotted against birth season (January/February/March) NAO index. Symbols differentiate different clusters defined by *k*-means cluster analysis of residual to the left-most set (black symbols).

**Fig. 13.**
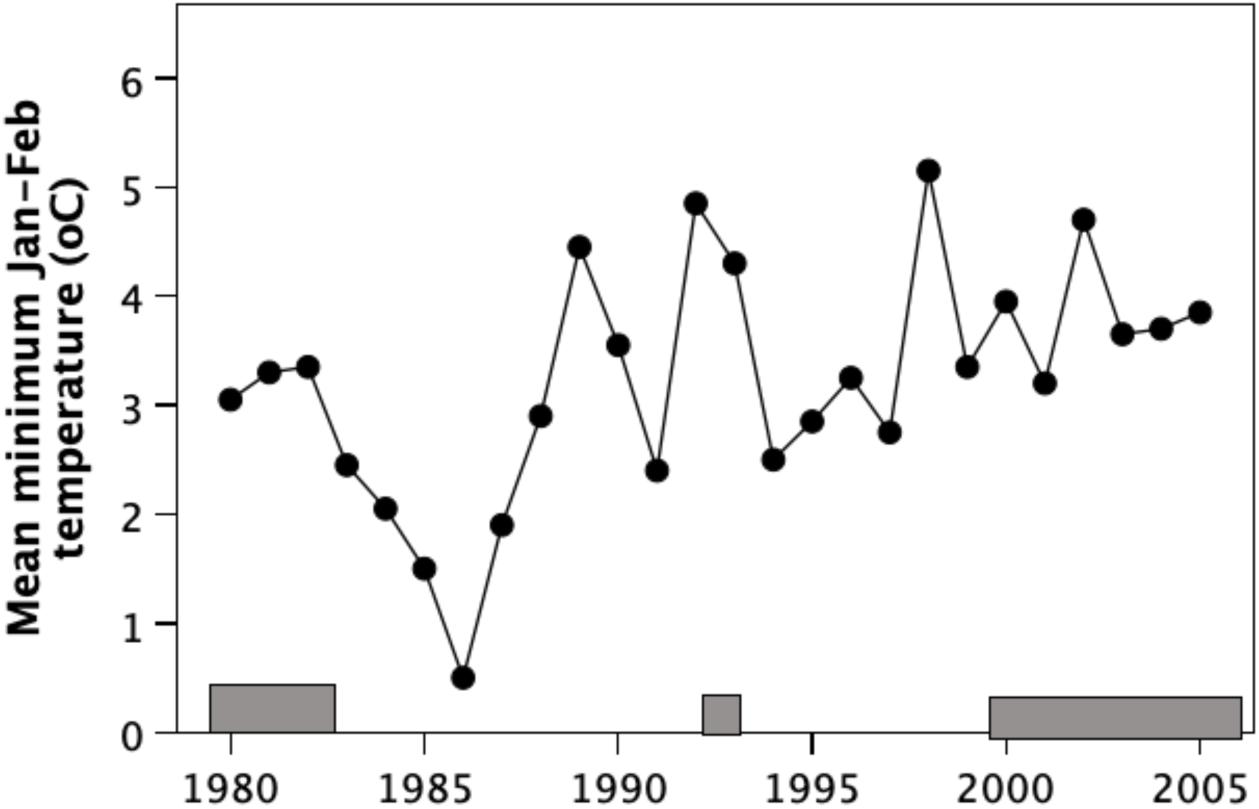
Mean minimum winter (January/February) temperature in each year. Study periods are demarcated as grey bars at the foot of the graph.

A backward stepwise multiple regression with growth rate as the dependent variable and population size in the preceeding year, cull intensity (% of population removed in previous autumn cull), NAO, rainfall in current and previous year, and NAO grade as dependent variables yields a significant regression in only the following variables:
GrowthRate_t_ = 1.692 − 0.001+Rain_t-1_ + 0.212*NAOwinter − 0.551+Grade
(overall model: F_3,9_=5.05, p=0.025; rainfall in t-1, t=3.51, p=0.007; NAO, t=-2.51, p=0.033; 3 grades, t=-3.89, p=0.004), where GrowthRate_t_ is the proportional increase in population size between year(t-1) and year(t) and Rain_t-1_ is the total rainfall (in mm) in the previous year.

Since the lack of any density-dependent effect seems surprising, I checked whether there was a density effect by regressing growth rate in each year directly on population size in the previous year. Although the relationship is negative (as might be expected), it is not significant (r^2^=0.090; t_11_=-1.04, p=0.160 1-tailed). I also reran this regression with populations size adjusted for cull intensity (%age of animals culled), but the results were the same. This suggests that environmental stresses had a much stronger effect than ecological competition between animals did. This is in line with the finding by Shi & Dunbar (2005) that conflict over food sources in this population are relatively rare (on average, one every 3.5 hours per individual female) even in late summer when forage quality is deteriorating. Moreover, such conflict was typically low key: in 86% of cases, the victim simply moved away to continue feeding elsewhere without involving any physical contact.

Since these analyses were run with NAO values for winter months (January-March), the period of peak kid vulnerability, I reran the analyses with NAO for the summer months (June/Juy/August) and for the year as a whole. However, these yielded less good or even non-significant fits, suggesting that it is the winter birth season conditions that are critical, presumably either through their impact on kid survival or indirectly via the cost of lactation and the effect this has on the females’ ability to survive the following summer. Note that mortality rates for winter-born kids do not differ between 1981 and 1982 despite very significant differences in weather conditions: 8 of 20 kids (40.0%) born between January and March died before 6 months of age in 1981 compared to 10 of 23 kids (43.5%) in 1982 (χ^2^=0.05, *p*>0.90), despite nearly twice as much rainfall in 1982. This suggests that the factor that most influences year-to-year variation in population size is not kid mortality as such but adult mortality. The relationship with winter NAO suggests that warmer late winter temperatures result in reduced adult survival. One explanation for this might be that warmer winters associated with higher rainfall result in the animals having to cope with near-freezing winter temperatures in pelts that are often waterlogged.

Although fecundity was somewhat higher in 1982 than 1981, the net growth rate for the population was significantly reduced in 1982 (R_0_=1.605 vs 1.062, respectively) despite a much higher birth rate because female mortality was much greater, especially among the prime breeding age females. Mean age at death for females who survived to 12 months of age was 9 years in 1981 but only 6 in 1982. As a result, the mid-life plateau in survivorship collapsed at about 6 years of age in 1982 compared to 9 years of age in 1981 (Fig. 14). This may have been due to the fact that, in 1982, late winter was much wetter than in 1981 (January-March rainfall: 433 vs 273 mm, respectively) and early summer much drier (April-July rainfall: 188mm vs 245mm), marking the beginning of a period of increasingly cold winters (Fig. 13). Irrespective of the causes of high mortality, Leslie matrix projections for population growth over the next 20 years indicates that the high mortality rates in 1982 would have resulted in a near-stationary population, whereas the other three years would have yielded very significant growth rates (Table 3).

**Fig. 14.**
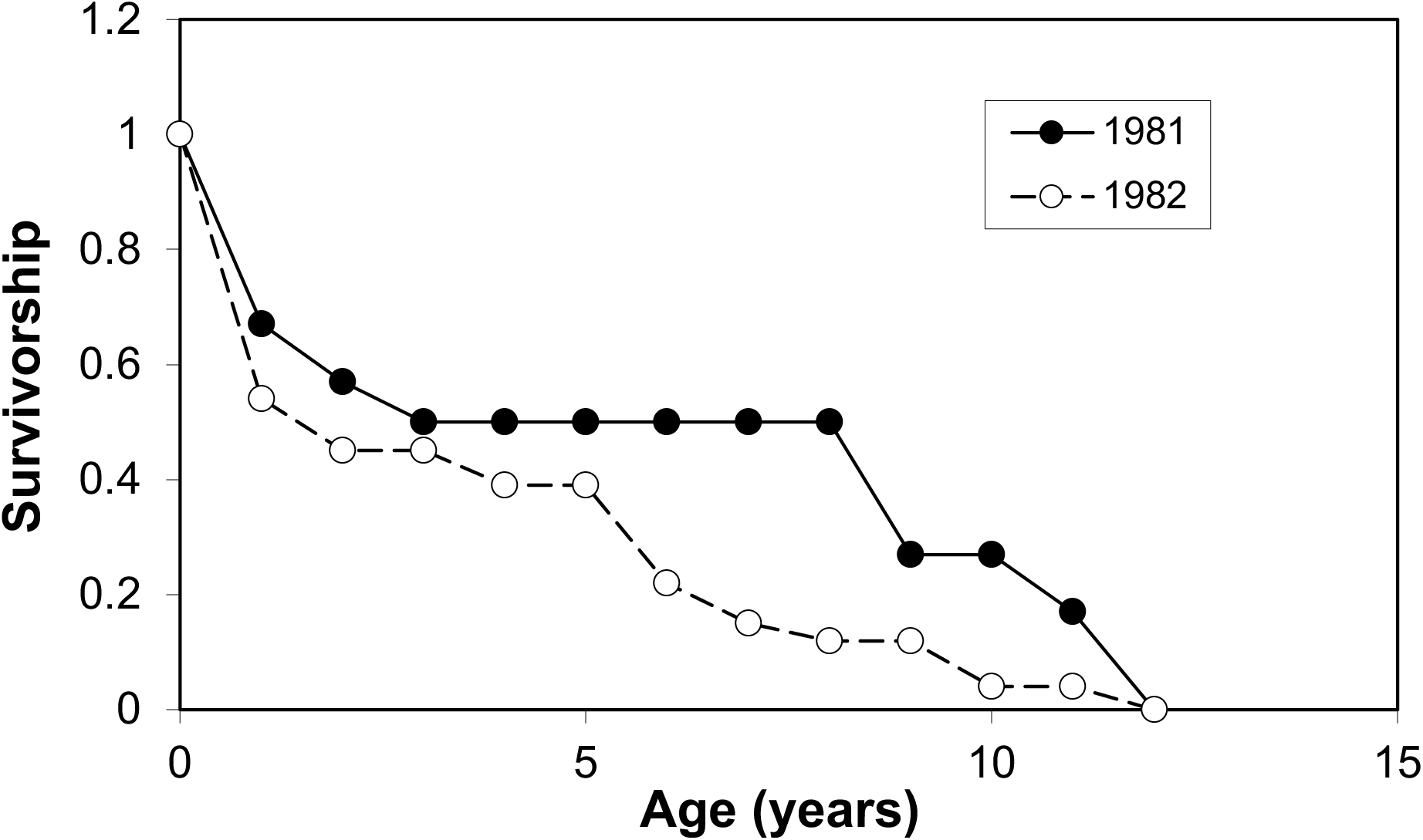
Age-specific survivorship curves for females during 1981 and 1982, averaged across the five sampled hefts. The data for 1982 were extrapolated to a full year using the data for equivalent months in 1981.

**Table 3.**
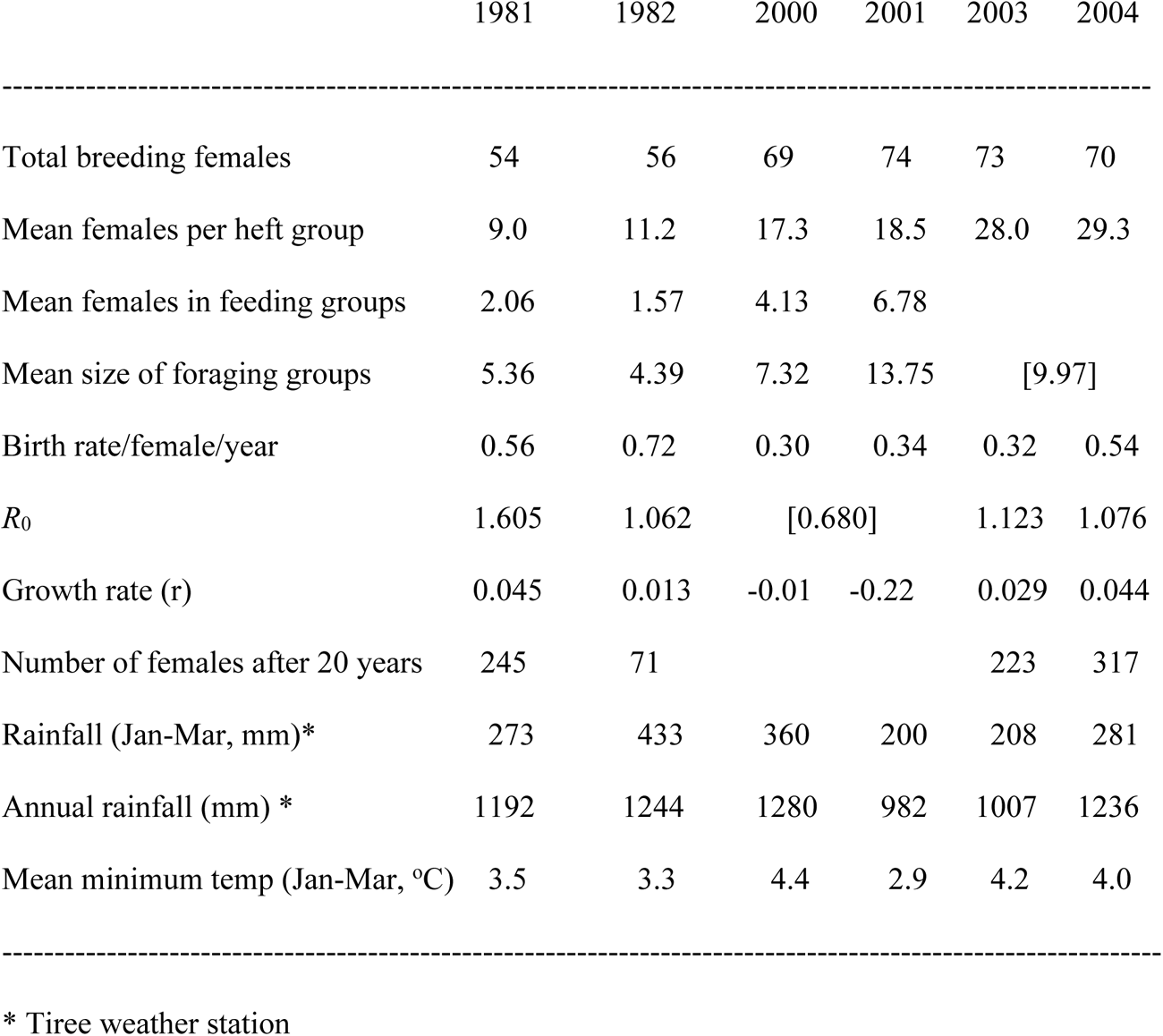
Demographic parameters and projections for different study years.

This climatically stressful period seems to have come to an abrupt end sometime in the later 1990s with the onset of a period of rapid population growth (Fig. 11) during which the main heft groups increased dramatically in size, culminating in the establishment of a new heft group occupying the low-lying grasslands of Harris Bay at the southern end of the study area sometime after 1993 (the last date we know for sure that they were not present in Harris). Note, however, that this period of rapid population growth was briefly brought to an abrupt halt by the onset of an unusually cold, dry phase in 2000 that resulted in negative population growth, before conditions improved again and growth rates became positive once more (Table 3).

This dramatic growth in the size of Harris/GnP/A’Bhrìdeanach hefts seems to coincide with the introduction of a culling programme for the red deer in the northern (Shellesder) block in 1992 (Fig. 15). Two points may be noted about deer numbers. First, the block on the north end of the goat study area (Shellesder/Sgorr Reidh) had very high numbers of deer, which, judging by the population’s stability was probably at carrying capacity for the local habitat – most likely because the next block to the north (Kilmory) had been a no-stalking zone from the 1970s onwards as part of the deer research project, and was exporting excess population to neighbouring blocks (Clutton-Brock et al. 2002). Second, although the southern blocks (GnP/Harris and Papadil/Dibidil) were also numerically quite stable, there is a marked downturn in their sizes from 1998 (coinciding with a switch in cull emphasis from the northern block to the southern blocks). Although overall goat numbers do not seem to have been affected by the 1992 cull in the northern block (Fig. 15), the neighbouring goat hefts (A’Brideanach, Sgorr Reidh and GnP) did increase dramatically during the ensuing years (Fig. 11). Conversely, while these hefts remained numerically stable from 2000 (Fig. 11), the population on the south side of the study area (the Papadil and Runcival hefts) did seem to increase dramatically following the decrease in the southern deer population (Fig. 15), resulting in the establishment of a new heft on the Harris Bay grasslands in the late 1990s (Fig. 11), leading to coinsiderable increase in the goat population by the end of the following decade (Fig. 15). By 2010, the goat population was estimated to be 357, double the typical size recorded during the 1960s-1990s.

**Fig. 15.**
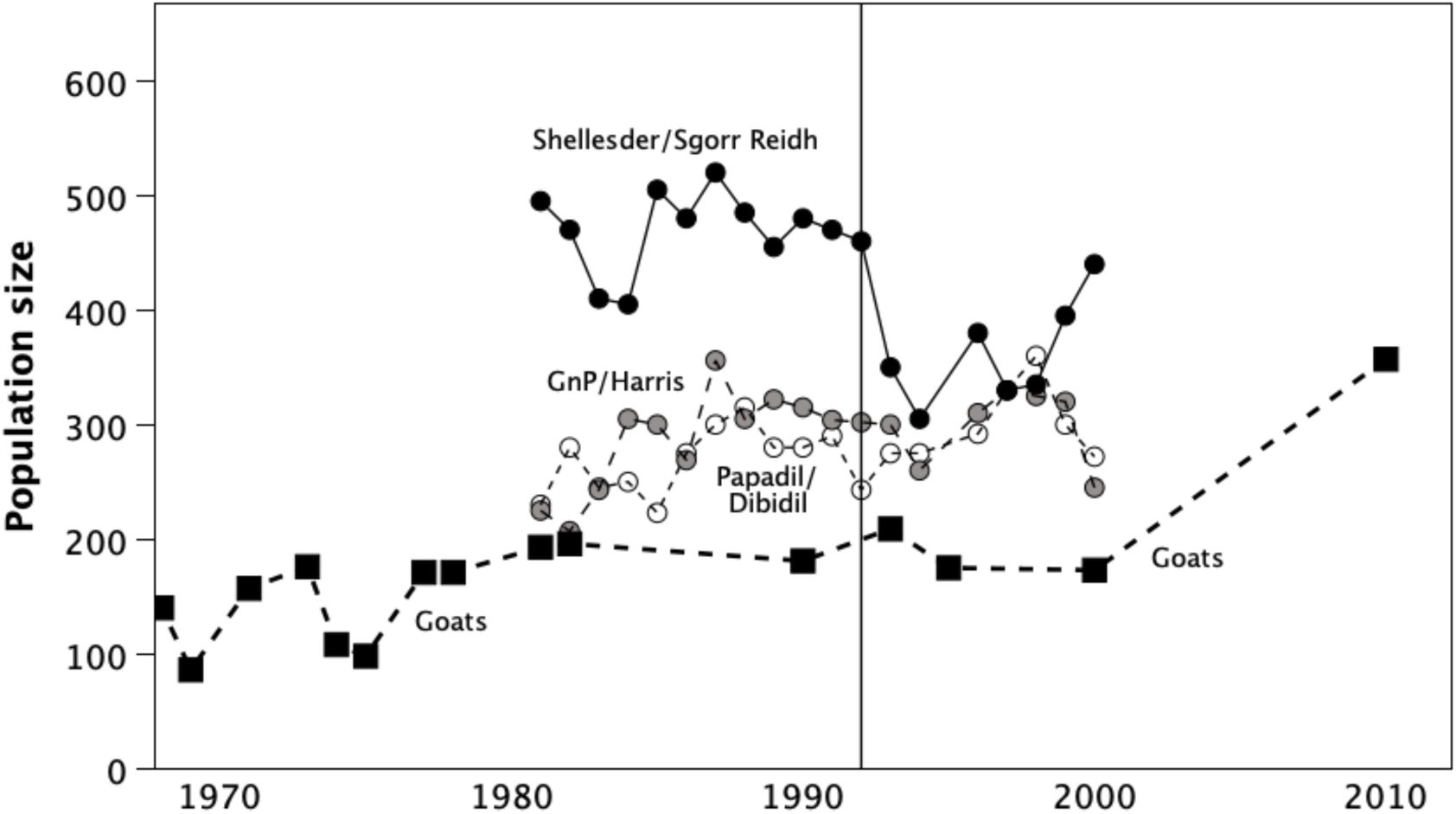
Population sizes of red deer (circles) in the three blocks corresponding to the goat study area (north to south: Shellesder/Sgorr Reidh, GnP/Harris, Papadil/Dibidil) and for the total goat population (squares) over time. The fine vertical line denotes the onset of the heavy deer cull in the northern (Shellesder) block in 1992. Sources: red deer: Clutton-Brock et al. (2002); goats: NatureScot goat censuses.

## Discussion

In contrast to populations of feral goats occupying many tropical and sub-tropical habitats, the Rùm goat population seems to have been in a state of quasi-equilibrium until the late 1990s, oscillating around a total population size of 100-200 animals. It seems likely that this was at least in part due to the stressful climatic conditions prevailing on these Hebridean islands. Rùm is close to the northern biogeographical limit for goats, as determined by the species’ ability to survive winter conditions (Dunbar & Shi 2013). The theoretical and actual northern limit is at the latitude of the Summer Isles, just 120 km north of Rùm. The importance of winter thermal conditions on both kid and adult female survival is symptomatic in this respect, since females’ ability to mitigate these thermal effects depends to a large extent on the availability of natural caves in which to shelter at night.

Density-dependence has been viewed as an important ecological factor influencing the population dynamics of the red deer population on Rùm (Clutton-Brock et al. 1987), as well as other north European grazing ungulates (Coulson et al. 2000). In the case of the Rùm goat population, there appears to be no density-dependent effect reflecting ecological competition, though it is possible that this is mainly a reflection of the small size of the population prior to the 2000s (Fig. 11). Analysis of feeding competition suggested that this is actually very low in the goats, and unlikely to have a significant impact on population dynamics (Shi 2002; Shi & Dunbar 2005). It seems likely that the goat population has been nowhere near its ecological ceiling.

Density-dependent effects can, however, arise from factors other than ecological competition. This can include the psycho-physiological stresses created by group-living (Dunbar & Shultz 2021) or as a result of males harassing females during the rut (Réale et al. 1998). There is some evidence, at least for the 1981-2 study cohort, that there was a significant negative effect of female heft group size on fecundity that parallels similar findings in other small, weakly social mammals (rodents, suids, carnivores) (Dunbar & Shultz 2021). In addition, the level of male harassment during the rut is very significant (Dunbar et al. 1990) and is often extremely disruptive for the target female’s foraging and rest. These possibilities require more detailed investigation beyond the scope of the present study.

On the other hand, there is strong circumstantial evidence for an effect of competition from the sympatric deer population. This may have been responsible for historically confining the goats to the island’s more rugged west coast. It seems likely (though the evidence is necessarily circumstantial) that it has been competition from the increasing deer population that has constrained the historical size of the goat population. However, it is difficult to differentiate between effects due to the reduction in deer numbers and those due to steadily rising winter temperatures that reflect the pace of global warming. Mean minimum winter temperatures rose by around 1.5°C over a 25-year period (Fig. 13), which would have translated into a significant 15% reduction in kid mortality (Fig. 10). It is clear that winter thermal conditions, when kids are vulnerable and females stressed to their limit by the costs of lactation, have a significant impact on mortality, and hence population dynamics. The steady relaxation of this thermal effect over time (Fig. 13) must inevitably have a significant long term effect on population size.

Although kid mortality at the population level seems to be mainly a function of ambient temperature during the first month or so of life, between-heft differences in kid mortality (in contrast to adult female mortality) seem to be related more directly to forage availability rather than nighttime shelters, suggesting that energy throughput may be a serious problem for lactating females once they have given birth. Our data identified heath habitat availability as the critical factor influencing kid survival. Gordon (1989b) noted that the dietetic selection of the Rùm goats was positively correlated with the biomass of dwarf shrubs (heaths) and negatively related to grass biomass (for a similar finding from southwest Scotland, see Bullock 1985). This may be one reason why the goats avoided Harris Bay for so long, since this is dominated by grasslands, with limited browse vegetation. One implication is that the new Harris heft may have experienced some constraints on its fertility, at least during its first few decades in a new habitat.

These findings point to the costs of lactation for females as the principal cause of heft differences in kid mortality, particularly given that most of this occurs within the first 2-3 months of life. Females stressed by low winter temperatures may be unable to obtain sufficient energy from their food to generate adequate milk supplies, especially if they live in poor quality habitats (i.e. those characterised by a high proportion of water-logged habitat and a low proportion of the drier heath habitats).

The low temperatures predominating during the late winter birth and lactation periods are probably the crucial issue here, because summer born kids suffer much lower mortality than winter-born kids. Hellawell (1991) also reported significant inter-heft variation in both fecundity and mortality rates in a Welsh population of feral goats, but was only able to consider altitude as a likely factor (in this case, mortality increased and fecundity decreased with altitude, implicitly suggesting that temperature may once again have been implicated). The fact that the birth season is in mid-winter contrasts starkly with the fact that, in the Mediterranean latitudes where the species evolved, goats give birth in spring (April-May), when foraging conditions would be at their best for lactation. This seems to be due to the fact that the rut is triggered by a specific declining daylength (D’occhio & Suttie 1992), and this occurs in late-summer on Rùm but in mid-autumn in the Mediterranean (Nicholson & Husband 1992; Papaioannou & Lovari 2023).

The importance of temperature in underpinning goat survival in habitats like Rùm is underlined by an analysis of the factors that influence the goats’ feeding time budgets: temperature turns out to be critical (Dunbar & Shi 2013). Temperature is also the key variable influencing tropical primates’ capacity to survive both in the habitats they currently occupy (Dunbar et al. 2009) and in the face of global climate warming (Korstjens et al. 2010, Lehmann et al. 2010).

The North Atlantic Oscillation index has been shown to be an important factor influencing the productivity in large mammals in northern Europe (Post et al. 1997, Forchhammer et al. 1998, Post & Stenseth 1999, Mysterud et al. 2001). High positive winter NAOs are associated with warm wet winters and, for coastal populations of deer at least, these are in turn associated with increased mean body mass and reduced fecundity. The population dynamics of the Rùm goats seem to respond in a similar way, with negative growth rates following winters when the NAO index was negative. That the relationship is explicitly with birth season NAO (and not NAO indices at other potentially important times of year) suggests that the principal factor lies with the difficulties that females have meeting their own energetic requirements during this period of the year as much as with kid survival *per se*. This also suggests that two rather different kinds of ecological processes are at work: population-wide effects related to largescale NAO-driven changes in late winter thermal conditions and a local-scale effect that is related to micro-climate (principally temperature) at the heft level. The latter mainly seems to reflect the availability of shelters (both in terms of caves and windbreaks) that ameliorate the impact of the larger scale effects.

The heft-scale effects can be expected to have a significant impact on population genetic structure if the various heft groups contribute differentially to the population over time. Some hefts may thus function in times of very adverse climatic conditions as source populations from which recolonisation of local areas subsequently occurs if hefts become locally extinct (as must be the case at Harris, since it separates the two halves of the population which must once have been connected). The interaction between cross- and within-subpopulation effects may thus be both significant and have important longterm consequences at the population level.

It seems that, until the later 1990s, the Rùm goat population was only just about able to hold its own under the prevailing climatic conditions (see also Dunbar & Shi 2013). However, rising winter temperatures combined with a reduction in deer numbers following the introduction of the deer cull in 1990 created conditions of ecological release for the goats and allowed numbers to increase steadily through the 2000s. Gordon & Illius (1989) found that female goats (but not the males) and red deer hinds had significant overlap in foraging patterns, suggesting that ecological competition between the two species is likely. More importantly, perhaps, Gordon et al. (1987) reported that goats tend to give way to the larger-bodied deer when they come into direct competition. This may explain why goats are mainly confined to the steeper hillsides and cliff faces on the west side of Rùm which deer tend to avoid (Gordon & Illius 1989).

The prominence of temperature as a regulating factor means that significant future changes in climate (such as that to be expected under climate warming) may have a very significant impact on the goats’ population dynamics. Under the climatic conditions prevailing in the 1980s, occasional severe winters (and especially the plunging winter temperatures during the early 1980s: Fig. 13) were sufficient to remove the excess population resulting from previous mild winters. With even modest levels of winter warming, this natural break might now be removed.

